# Metabolic Engineering Of *Lactococcus Lactis* For The Production Of Heparosan

**DOI:** 10.1101/2022.12.28.522110

**Authors:** Siddharth Guhan, Naveen Raj, Senthilkumar Sivaprakasam, Pandeeswari Jeeva

## Abstract

Heparosan is a precursor molecule for the widely used anticoagulant heparin, which also has other uses such as certain drug delivery applications and as a scaffold for tissue engineering in biomaterials. Traditionally, pathogenic bacteria such as *E*.*Coli* have been used as a host to produce heparosan as an alternative to animal and chemoenzymatic synthesis. Using GRAS status organisms like *Lactococcus Lactis* as the host for production of heparosan provides a safe alternative as well as being a well-established organism for genetic manipulation and reengineering. In this study, a functional heparosan synthesis pathway was successfully expressed in *Lactococcus Lactis* by the expression of *E*.*coli* K5 genes KfiA and KfiC, along with the overexpression of ugd, glmu and pgma genes present natively in the host organism. The genes were activated using the tightly controlled NICE expression system. The genes were cloned into plasmid p8148 and transformed into two strains, *Lactococcus Lactis* NZ9000 and *Lactococcus Lactis* NZ9020, totaling six different recombinant strains were created using these two hosts and various combinations of the heterologous genes. The recombinant *Lactococcus Lactis* SH6 strain, expressing the genes *ugd-KfiA-KfiC-*pgma yielded a maximum concentration of 754 mg/l in batch bioreactor experiments and the titer was increased to 1263 mg/l in fed-batch fermentation. NMR imaging successfully determined that the structure of the product derived from *Lactococcus Lactis* was indeed similar to *E*.*coli* heparosan. The molecular weight of heparosan varied from 10-20 KDa, indicating its potential use for chemoenzymatic heparin biosynthesis.

---

The current demand for antithrombotic agents is becoming extensive, especially for cardiovascular disorders like arterial/venous thrombosis, which accounts for a substantial mortality rate (Singh et al., 2019). Heparin is the most prominent parenteral or direct-acting anticoagulant exploited to overcome the natural propensity for blood to clot and prevent thrombosis (Baik et al., 2012). Anticoagulation occurs when heparin binds with antithrombin III (ATIII), a serine protease inhibitor (serpin) which undergoes a conformational change, and it gets activated as an inhibitor of thrombin and other serine proteases in the coagulation cascade (Linhardt, 2003). Commercial availability of heparin is extracted from the mucus membrane of the pig intestine and ox lungs. Animal-derived heparin comprises process impurities, viruses, and prions (Linhardt, 2003). The presence of over-sulphated chondroitin sulphate led to serious health issues for US people in 2008 (Kishimoto et al., 2008). As a result, US FDA has improved the pharmacopeial monographs to reduce the likelihood of this crisis in the future (Paluck et al., 2016). Because of this, the demand for heparin has risen a lot, and to meet the requirements of the same several methods like chemical synthesis and chemoenzymatic methods have been developed. The chemical synthesis of heparin is a highly tedious and complicated process to synthesize oligosaccharides. Fondaparinux, a chemically synthesized heparin in the market, requires around 60 steps for synthesis, with meager yield and high cost (Linhardt & Liu, 2012). The production of biosynthetic heparin through the in-vitro chemoenzymatic method offers a safer alternative approach, but it requires heparosan, an un-sulphated and unepimerized heparin backbone, as a precursor molecule for both heparin as well as heparin sulphate synthesis (Xu et al., 2011) (Zhang et al., 2012). In addition to its antithrombotic activity, heparin, as well as heparin sulphate, is used in the treatment of cancer to prevent metastasis because of its interaction with the growth factors (Yip et al., 2006) and in the prevention of virus infections (Bishop et al., 2007).

Heparosan is a linear glycosaminoglycan (GAG) consisting of repeated disaccharide units of D-glucuronic acid (GlcUA) and N-acetyl-D-glucosamine (GlcNAc) linked through α-1,4 and β-1,4 glycosidic bonds. Owing to the presence of the carboxylate group in glucuronic acid, heparosan is having highly hydrophilic and anionic property which makes it suitable for conjugating them with the liposome for drug delivery and dissolution purposes (Lane et al., 2017). Heparosan is beneficial for medication delivery, dermal fillers, and biomaterials which possess qualities including moisture retention capacity, strong non-immunogenicity, and biocompatibility (Linhardt & Liu, 2012). Microorganisms that naturally produce heparosan will secrete it to invade the host immune system and serve as a biofilm for their survival.

Natural producers of heparosan, namely *E. coli* K5 and *Pasteurella multocida*, are highly pathogenic and cause urinary tract infections. Thus, it would be advantageous to look for an alternative microbial source for producing heparosan to avoid the potential crisis (Zhang et al., 2012). The molecular weight of the heparosan is one of the essential product qualities of heparosan which is having a market value in the pharmaceutical industry. Low molecular weight heparosan was found to have effective anticoagulant/ antithrombotic activity as well as extended bioavailability (Linhardt, 2003). *E. coli* Nissle 1917 is a probiotic and natural producer of heparosan, produces high molecular weight heparosan of 68 kDa, and bimodal in distribution. Production of unimodal distribution of heparosan is crucial for purification and applications since it reduces the number of steps in downstream processing (Hu et al., 2022a).

Microbial production of heparosan from a non-pathogenic source is attaining higher importance to avoid any contamination crisis and make purification much easier (Hu et al., 2022b). Heterologous expression of glycosyl transferase enzyme-producing genes from the source of *E. coli* K5 and *P. multocida* to produce heparosan in non-pathogenic microorganisms is considered a safer alternative for human consumption. Even though *E. coli* is the most used prokaryotic system for gene expression, the significant bottlenecks associated with them are the formation of endotoxins due to the presence of lipopolysaccharide in its outer membrane and the formation of inclusion bodies which leads to misfolding of proteins and biologically unfunctional ones (Savvas C. Makrides & Gerhard Hannig, 1998). Natural producer of heparosan, *E. coli* K5 will produce low molecular weight heparosan which acts as a precursor for heparin. Therefore, to achieve the same expression in the *L. lactis*, glycosyl transferase genes from *E. coli* K5 were chosen (Ly et al., 2011)

Tremendous advances have been made in unraveling the genetics and molecular biology of these economically significant microorganisms. This wealth of knowledge and experience has led to the use of *lactococci* far beyond their original role in food preservation and production. *Lactococcus lactis* is the most widely used and Generally Recognized as safe (GRAS) microorganism in the food and dairy industries. The growth of *L. lactis* is very rapid (lower doubling time), which makes it easier to achieve high cell density also aeration is not essential for its growth, suitable for large-scale fermentation (King et al., 2015a). Many tools and techniques have been developed to genetically engineer *L. lactis*, such as plasmid cloning vectors, various (inducible) gene expression vectors, and methodologies to introduce any mutation in specified regions of the *L. lactis* genome. *L. lactis* does not produce endotoxins and inclusion bodies in contrast to *E. coli* K5, which facilitates the use of value-added products expressed by *L. lactis* in clinical and food applications (Elmarzugi et al., 2010). Additionally, the number of steps required in the downstream processing is reduced by the absence of endotoxins. This has led to its widespread application in the recombinant synthesis of numerous therapeutic enzymes and vitamins. In *L. lactis* due to the presence of a tight nisin A promoter, there is no selection pressure on the maintenance of the plasmids, resulting in plasmid stability and the great reproducibility of expression trials (King et al., 2015b). Nisin control-based expression system (NICE) is a very tightly controlled gene expression in *L. lactis* NZ9000, leading to undetectable protein expression in the uninduced state, which serves as an advantage during the plasmid maintenance in the maintenance host. NICE is based on a two-component regulatory system, namely *nisR* and *nisK*, which will be getting activated when the nisin binds to the receptor and the P_nisA_ promoter got activated (Kleerebezem et al., 1997). Furthermore, the hyaluronic acid structural analog of heparosan was successfully cloned and expressed in *L. lactis* (Chien & Lee, 2007). These characteristics endorse the selection of *L. lactis* as a cloning host to produce heparosan. The performance of the recombinant strains was evaluated in both shake flasks and bioreactor. The structural and molecular weight characterization of heparosan was elucidated. This study revealed a secure alternate method to synthesize heparosan for the production of bioengineered heparin.

## Materials and methods

### Media components

M17 media components (2.5 g/L casein hydrolysate, 2.5 g/L animal digest of peptone, 5 g/L of soya peptone, 2.5 g/L yeast extract, 5 g/L beef extract, 0.5 g/L ascorbic acid, 0.25% MgSO_4_.7H_2_O, 27.9 g/L hydrated sodium-β-glycerophosphate) glucose, and cetyltrimethylammonium bromide (CTAB) were purchased from HiMedia Laboratories (India). Isopropyl alcohol, sulfuric acid, and sodium nitrate were purchased from Merck (USA). The GenElute Bacterial Genomic Kit, chloramphenicol, and nisin were obtained from Sigma-Aldrich (USA). QIAprep Spin Miniprep kit for plasmid isolation, QIAquick™Gel Extraction kit, and the QIAquick™ PCR purification kit were purchased from Qiagen (Germany). The enzymes NcoI, KpnI, PmlI, Phusion™DNA Polymerase, and T4 DNA ligase were purchased from New England Biolabs (USA).

### Construction of recombinant L. *lactis* clones for expressing *ugd, kfiA, kfiC, glmu*, and *pgmA* genes

The recombinant plasmids and bacterial strains used in this study are given in Table 1. The genomic DNA of *L. lactis* NZ9000 was isolated using GenElute™ Bacterial Genomic DNA Kit. Using the Phusion™ High-Fidelity DNA Polymerase, the *ugd, pgma*, and *glmu* genes were amplified from *L. lactis* NZ9000 genomic DNA. The protein encoding sequence of the *kfiA* and *kfiC* genes were retrieved from *E. coli K5* and codon optimization was done (GenScript). The primers used for amplifying different gene fragments and the associated restriction sites are shown in Table 3. The amplified gene fragments were cloned into pNZ8148 by the conventional cloning method. *E. coli* MC1061 was used as a cloning host to construct recombinant plasmids expressing the genes *kfiA* and *kfiC*. The resulting recombinant plasmids were transformed into *L. lactis*.

**Table 1:** Recombinant plasmids and strains used in this study for the heparosan production in *L. lactis*

| Strain/plasmid | Description/characteristic | Source/Reference |
| --- | --- | --- |
| Strain |  |  |
| *E. coli* K5 | Serotype O10:K5:H4 | BEI Resources, NIAID, NIH |
| *E. coli* MC1061 | *araD139*, $\Delta$ (*ara*, *leu*)7697, *ΔlacX74*, *galU*<sup>-</sup>, *galK*<sup>-</sup>, *hsr*<sup>-</sup>, *hsm*<sup>+</sup>, *strA* | National Institute of Immunology, New Delhi. |
| *L.lactis* NZ9000 | MG1363 strain containing *nisRK* genes integrated into *pepN* locus, plasmid-free | NIZO Netherlands |
| *L.lactis* NZ9020 | *ΔldhX* and *ΔldhB* knock out of NZ9000 with *Tet<sup>R</sup> Ery<sup>R</sup>* | NIZO Netherlands |
| *L.lactis* SH2A | *L. lactis* NZ9000 expressing *kfiA-kfiC-ugd* gene in plasmid pNZ8148 | This study |
| *L.lactis* SH4A | *L. lactis* NZ9000 expressing *kfiA-kfiC-glmu* gene in plasmid pNZ8148 | This study |
| *L.lactis* SH6A | *L. lactis* NZ9000 expressing *kfiA-kfiC-ugd-pgma* gene in plasmid pNZ8148 | This study |
| *L.lactis* SH2 | *L. lactis* NZ9020 expressing *kfiA-kfiC-ugd* gene in plasmid pNZ8148 | This study |
| *L.lactis* SH4 | *L. lactis* NZ9020 expressing *kfiA-kfiC-glmu* gene in plasmid pNZ8148 | This study |
| *L.lactis* SH6 | *L. lactis* NZ9020 expressing *kfiA-kfiC-ugd-pgma* gene in plasmid pNZ8148 | This study |
| Plasmid | Nisin inducible vector with *Cm<sup>R</sup>* | NIZO Netherlands |
| pNZ8148 |  |  |
| pSH2 | pNZ8148 with inserted *kfiAC-ugd* genes | This study |
| pSH4 | pNZ8148 with inserted *kfiAC*-<br>*pgma* genes | This study |
| pSH6 | pNZ8148 with inserted *kfiAC-ugd*-<br>*pgma* genes | This study |

The PCR conditions for the amplification of the gene fragments are listed in Table 2. The amplicons were then purified using a QIA quick™ PCR purification kit. The expression of heparosan was controlled by placing them under the influence of a strong promoter, namely *P*_*nisA*_. Nisin inducible plasmid, pNZ8148, was used to construct recombinant plasmids pSH2 (containing *kfiA, kfiC*, and *ugd* genes), pSH4 (containing *kfiA, kfiC*, and *glmu* genes), and pSH6 (containing *kfiA, kfiC, ugd* and *pgmA* genes). The ligated products were transformed into ultra-competent *E. coli* MC1061 and plated on Luria-Bertani agar plates containing chloramphenicol (10 μg/mL) and streptomycin (10 μg/mL). The final plasmid construction map of all the plasmids is shown in Figure1. The positive recombinants were screened using colony PCR. Double restriction digestion and DNA sequencing were used to confirm the same. The resulting plasmid-containing cells were transformed into the host *L. lactis NZ9000* and *L. lactis NZ9020* by electroporation.

**Table 2:** PCR conditions used for the amplification of the genes and their corresponding temperature

| Steps | Time(s) | Temperature |  |
| --- | --- | --- | --- |
| Initial denaturation | 120 | 98 °C |  |
| Denaturation | 30 | 98 °C |  |
| Annealing | 45 | *kfi AC* | 53 °C |
|  |  | *ugd* | 52 °C |
|  |  | *glmu* | 52 °C |
| Extension | 45 | 72 °C |  |
| Final extension | 180 | 72 °C |  |

**Table 3:**
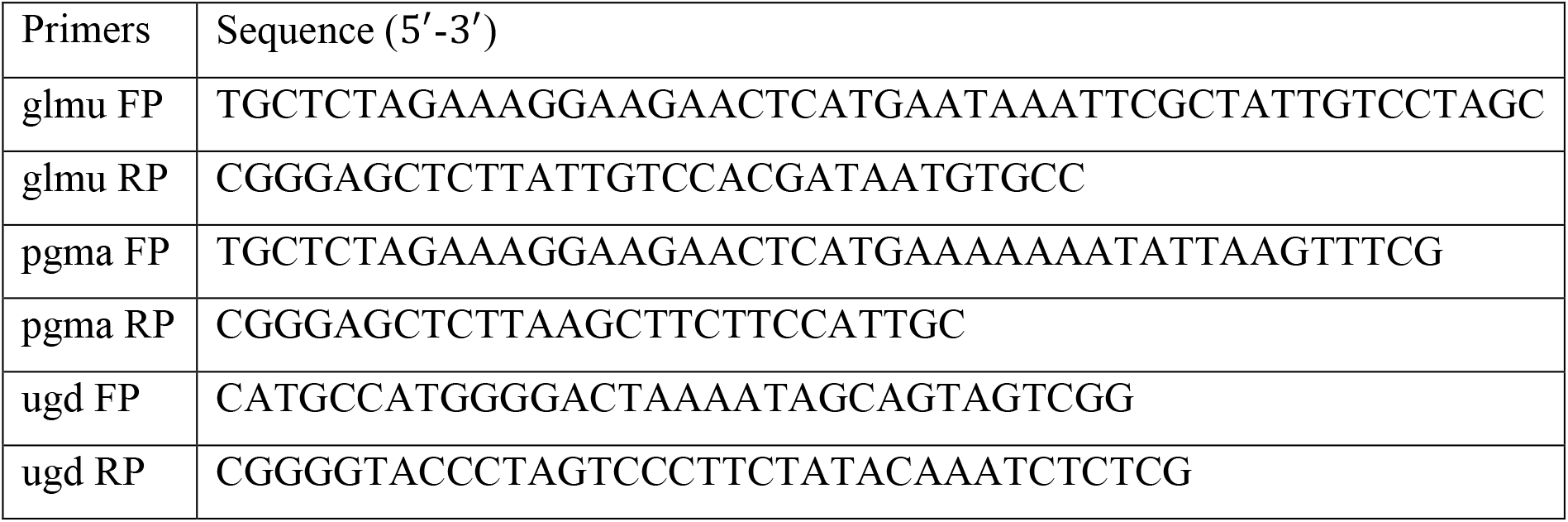
Forward primer (FP) and reverse primer (RP) sequence used for cloning *glmu, pgma*, and *ugd* genes into plasmid pSH2, pSH4, and pSH6.

**Figure 1:**
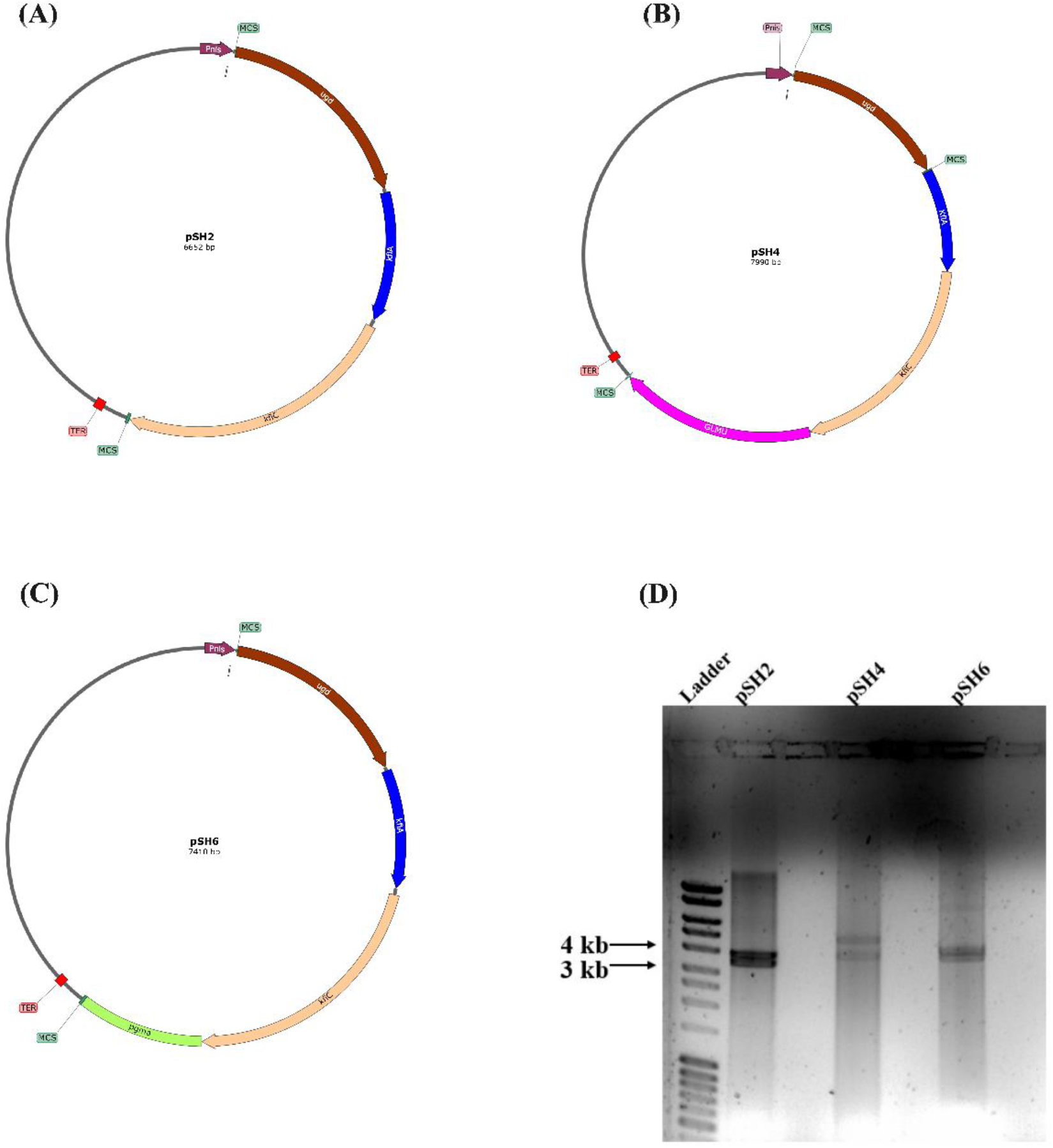
Plasmid map construction of (A) pSH2 (B) pSH4 and (C) pSH6. (D) Clone confirmation by double restriction and digestion of recombinant plasmids pSH2, pSH4, and pSH6.

### Clone comparison and selection

Table 1 shows the various gene combinations used to construct the six clones in the two strains, NZ9000 and NZ9020. In the shake flask, clone screening was performed to determine which of the six clones is the most efficient in terms of product titer and yield. Preculture was prepared by using a single colony from the freshly prepared plate and 2% (v/v) of preculture was inoculated into the M17 media containing the glucose (15 g/L), and chloramphenicol (10 μg/mL) was added for the strain NZ9000. For the strain, NZ9020 M17 media containing glucose (15 g/L), chloramphenicol (10 μg/mL), tetracycline (2 μg/mL), and erythromycin (10 μg/mL) were used. The culture was grown at 30°C in static and shaking conditions at 150 rpm for 26 hrs. Induction was done using nisin (4 ng/mL) when the cell reached OD_600_ around 0.4-0.6. Samples were withdrawn periodically every 2 hrs, and the biomass, product, glucose, and lactate concentration, and resulting process parameters, namely specific growth rate, substrate uptake efficiency, yield coefficient, specific product formation rates, and yield coefficients, were calculated.

### Bioreactor fermentation experiments

The recombinant *L. lactis* SH6 was grown in the M17 media containing glucose in a shake flask for 12 hrs which serves as inoculum for the bioreactor (Biojenik engineering, India) of working volume 1.7 L. Modified M17 media (1.2 L) with the following composition was autoclaved ex-situ: yeast extract-5 g/L, brain heart infusion-5 g/L, MgSO_4_·7H2O-0.5 g/L, ascorbic acid-0.5 g/L, KH_2_PO4-0.5 g/L, and K_2_HPO_4_-1.5 g/L, autoclaved glucose (30 g/L), chloramphenicol (10 μg/mL), tetracycline (2 μg/mL), and erythromycin (10 μg/mL) were added. The temperature was maintained at 30°C, and pH was controlled at 7 by using 4M KOH. The agitation speed was maintained at around 200 to 400 rpm. The reactor was operated under two conditions: low aeration having less than 5% Dissolved Oxygen (DO) with an aeration rate of 0.5-0.6 vvm and high aeration having 15% DO with an aeration rate of 1-1.2 vvm maintained throughout the process. Fed-batch fermentation was also carried out in the 3L bioreactor (Biojenik engineering, India) for the recombinant *L. lactis* SH6 strain under high aerated conditions. Induction was done around the 8^th^ hr using nisin (4 ng/mL) when it reached the OD_600_ around 0.4-0.6. Glucose of inlet feed concentration 300 g/L was added around 20^th^ hr under aseptic conditions when the glucose was exhausted in the reactor. Samples were regularly withdrawn from the reactor to analyze biomass, product, glucose, lactate, and ethanol concentration.

### Analytical techniques

#### Determination of Biomass concentration

Biomass concentration was determined based on absorbance measurement at 600nm. The cells were harvested and centrifuged for 10 minutes at 13,000 rpm, and the cell pellet was resuspended in millipore water. After diluting the samples to acquire various Optical Density (OD) values ranging from 0.2 to 1, the samples were lyophilized. The resulting cell’s dry weight was determined, and a correlation between the OD and Dry Cell Weight (DCW) was created.

#### Heparosan concentration and molecular weight determination

Heparosan was quantified based on the CTAB assay (Oueslati et al., 2014). The collected samples were centrifuged at 13,000 rpm for 10mins, then resuspended in the Tris-EDTA buffer and ultrasonicated, followed by centrifugation to remove the cellular debris. The resultant supernatant was processed for the CTAB assay. High-performance size exclusion chromatography (HPSEC) equipped with a RI detector (Shimadzu, Kyoto, Japan) was used to determine the molecular weight distribution of heparosan. LabSolutions software (Shimadzu, Kyoto, Japan) was used to acquire and process the chromatogram data. Heparosan sample obtained from the fed-batch process after purification was filtered through a 0.45 μm filter and loaded into the polysep GFC-P-6000 column. A mobile phase of 0.1M NaNO_3_ was used at a flow rate of 0.5 mL/min, and the column temperature was maintained at 40°C. Dextran standards of different molecular weights 10 kDa, 20 kDa, 40 kDa, 70 kDa, and 100 kDa (Sigma Aldrich, MO, USA) were used to construct the calibration plot.

#### Quantification of glucose and lactate concentration

Glucose, lactate, and other metabolites concentration in the cell supernatant were measured using Ion exchange chromatography Phemomenex Aminex HPX-87H column and Phenomenex guard column fitted to Shimadzu HPLC-RID detector system were used for the analysis. A mobile phase of 5 mM H_2_SO_4_ was used, and the isocratic elution method was followed with a flow rate of 0.6 mL/min. Glucose concentration was determined by using the RID detector system. Lactate and other metabolites concentrations were determined by using UV at 210nm.

#### Heparosan purification

Aqueous two-phase extraction (ATPS) and dialysis were used for the purification of heparosan. The fermentation broth from the fed-batch process was centrifuged at 13,000 rpm for 10 mins, and it was resuspended in the lysis buffer (1 g/L Lysozyme, 0.5 mM EDTA, and Tris-HCl pH 7.5 and incubated for 2hrs. The resulting cell lysate was autoclaved at 121 °C for 21 mins (Nehru et al., 2020). The resulting supernatant was subjected to aqueous two-phase extraction using PEG 8000 and potassium phosphate (Rajendran et al., 2016). It was collected and processed for dialysis using the dialysis bag having the membrane cut off of 5 kDa against deionized water for 6 hrs, and the sample was lyophilized.

#### FTIR and NMR analysis

The purified heparosan sample was lyophilized, and both the crude sample and the purified sample were subjected to an FTIR spectrometer (Perkin Elmer, USA) with a scan range of 450– 4500 cm^−1^. H^1^ and C^13^ experiments were performed for the purified heparosan using Bruker Ascend 600 MHz NMR spectrometer (Bruker, MA, USA) with TOPSPIN data acquisition software (Bruker). The sample was dissolved in 400 μL of deuterated water (Sigma Aldrich, MO, USA) and lyophilized to facilitate hydrogen-deuterium transfer. The sample was then dissolved in 500 μL deuterated water and transferred to the 5 mm standard NMR tubes. ^1^H spectrum was acquired in water suppression mode for 128 scans. ^13^C spectrum was obtained for 6000 scans at 298 K.

## Results and Discussion

### Construction of recombinant *L. lactis*

The recombinant *L. lactis* strains were made by introducing the plasmids pSH2, pSH4, and pSH6 into the strains NZ9000 and NZ9020 to produce six different clones SH2, SH2A, SH4, SH4A, SH6, SH6A. The same was verified by using the double restriction enzyme cleavage and the gene sequencing shown in Figure 1. Enhanced heparosan production appears as a result of maintaining high intracellular precursor concentrations. Hence, precursors of heparosan, glucuronic acid, and N-acetyl glucosamine production should be increased (Badle et al., 2014a). To improve the intracellular precursor concentration, overexpression of genes *pgma, glmu* and *ugd* were done. In *L. lactis*, the expression of the *ugd* gene, which is involved in the production of glucuronic acid, is tightly controlled transcriptionally, serving as a rate-limiting step (Prasad et al., 2012). The generation of glucuronic acid was also minimal because the metabolites involved in its synthesis are also involved in synthesizing the peptidoglycans found in cell walls (Zhang et al., 2012). Consequently, the *ugd* gene was cloned in all three plasmids (pSH2, pSH4, and pSH6) to enhance the glucuronic acid concentration inside the cell. The *glmu* gene was overexpressed in the plasmid pSH4 to enhance the intracellular precursor N-Acetyl Glucosamine concentration. To elongate the polymer chain length of heparosan, the enzymes *kfiA* and *kfiC* are essential. Heparosan production in *L. lactis* is facilitated by the concerted action of the genes *KfiA* and *KfiC* and the formation of the *kfiA-kfiC* enzyme complex (Hu et al., 2022b). Thus, they were cloned in all three plasmids (pSH2, pSH4, and pSH6). One of the common intermediary metabolites involved in the production of both glucuronic acid and cell wall peptidoglycan is glucose-1-phosphate whose production was increased by overexpressing the gene *pgma* in plasmid pSH6, which is responsible for converting glucose-6-phosphate into glucose-1-phosphate. The engineered metabolic pathway in recombinant *L. lactis* for heparosan production was depicted in Figure 3.

### Clone selection

All six different clones were cultured in the flask under static and shaking conditions to determine which of the six clones was best in product titre, yield coefficient, substrate uptake efficiency, and production formation rate. The carbon source that *L. lactis* consumes is used to produce biomass, maintenance energy, and product formation. The product yield, biomass concentration, and yield coefficient parameters obtained in both static and shaking conditions are shown in Table 4. The product titre of the SH4 clone was found to be a maximum of 0.33 gL^-1^ at the static conditions. This might be due to overexpressing of both the genes, namely *ugd* and *glmu* which are involved in the production of both intracellular precursors N-Acetyl glucosamine and UDP-Glucuronic acid for heparosan production (Badle et al., 2014b). Even though the product titre was found to be a maximum for the SH4 clone under static conditions, it also produces the primary by-product of lactic acid of 0.457 gL^-1,^ which increases the cost of downstream processing. The yield coefficient of the product to the substrate Y_p/s_ and the specific product formation rate q_p_ for the SH4 clone was also low at 0.054 gg^-1^ and 0.029 gg^-1^hr^-1^. This indicates that most carbon flux is passing for by-products like lactic acid production. The same case was observed with the SH2 clone producing heparosan of 0.298 gL^-1^ and lactic acid of 0.817 gL^-1^ under static conditions with low Y_p/s_ and specific product formation rate q_p_ of 0.045 gg^-1^ and 0.028 gg^-1^hr^-1^. The lowest product titre was observed in the SH2 clone, possibly due to an imbalance of the intracellular precursor. To produce heparosan, the intracellular precursor concentration must be balanced; if one precursor is produced in excess compared to another, the glycosyl transferase activity will be terminated, which will halt the formation of heparosan (Badle et al., 2014a). Cloning the gene for glucuronic acid production, *ugd*, onto plasmid pSH2 could increase the concentration of glucuronic acid relatively excess than the N-Acetyl Glucosamine which might halt the glycosyl transferase activity resulting in lower heparosan production. It was also found that lactate production was found to be relatively higher for *L. lactis* NZ9000 (SH2A, SH4A, SH6A) than the *L. lactis* NZ9020 (SH2, SH4, SH6).

**Table 4:** Comparison of specific growth rate, product formation rate, substrate uptake rate, yield coefficient of biomass, product, lactic acid on the substrate, product titre, and lactic acid titre for the six different clones under static and shaking conditions.

| Clone | | $\mu$<br>(hr<sup>-1</sup>) | $q_p$<br>(gg<sup>-1</sup>hr<sup>-1</sup>) | $q_s$<br>(gg<sup>-1</sup>hr<sup>-1</sup>) | $Y_{X/S}$<br>(gg<sup>-1</sup>) | $Y_{P/S}$<br>(gg<sup>-1</sup>) | $Y_{LA/S}$<br>(gg<sup>-1</sup>) | $Y_{P/x}$<br>(gg<sup>-1</sup>) | Heparosan<br>(gL<sup>-1</sup>) | Lactate<br>(gL<sup>-1</sup>) |
| --- | --- | --- | --- | --- | --- | --- | --- | --- | --- | --- |
| SH2 | Static | 0.214 | 0.035 | 0.335 | 0.615 | 0.108 | 0.137 | 0.161 | 0.208 | 0.73 |
|  | Shake | 0.529 | 0.0058 | 0.729 | 0.701 | 0.008 | 0.17 | 0.017 | 0.155 | 1.15 |
| SH2A | Static | 0.418 | 0.028 | 0.587 | 0.752 | 0.045 | 0.151 | 0.071 | 0.298 | 0.817 |
|  | Shake | 0.583 | 0.187 | 1.02 | 0.509 | 0.23 | 0.25 | 0.298 | 0.19 | 1.45 |
| SH4 | Static | 0.476 | 0.029 | 0.682 | 0.677 | 0.054 | 0.149 | 0.087 | 0.33 | 0.457 |
|  | Shake | 0.294 | 0.103 | 0.592 | 0.477 | 0.172 | 0.202 | 0.325 | 0.141 | 0.75 |
| SH4A | Static | 0.155 | 0.034 | 0.437 | 0.363 | 0.1 | 0.472 | 0.241 | 0.244 | 0.622 |
|  | Shake | 0.468 | 0.018 | 1.11 | 0.44 | 0.018 | 0.482 | 0.035 | 0.187 | 1.12 |
| SH6 | Static | 0.162 | 0.059 | 0.435 | 0.579 | 0.197 | 0.003 | 0.368 | 0.262 | 0.036 |
|  | Shake | 0.349 | 0.127 | 0.524 | 0.725 | 0.182 | 0.003 | 0.358 | 0.201 | 0.044 |
| SH6A | Static | 0.223 | 0.018 | 0.47 | 0.396 | 0.04 | 0.478 | 0.087 | 0.119 | 0.778 |
|  | Shake | 0.322 | 0.0025 | 0.588 | 0.55 | 0.046 | 0.309 | 0.008 | 0.233 | 0.683 |

**Table 5:**
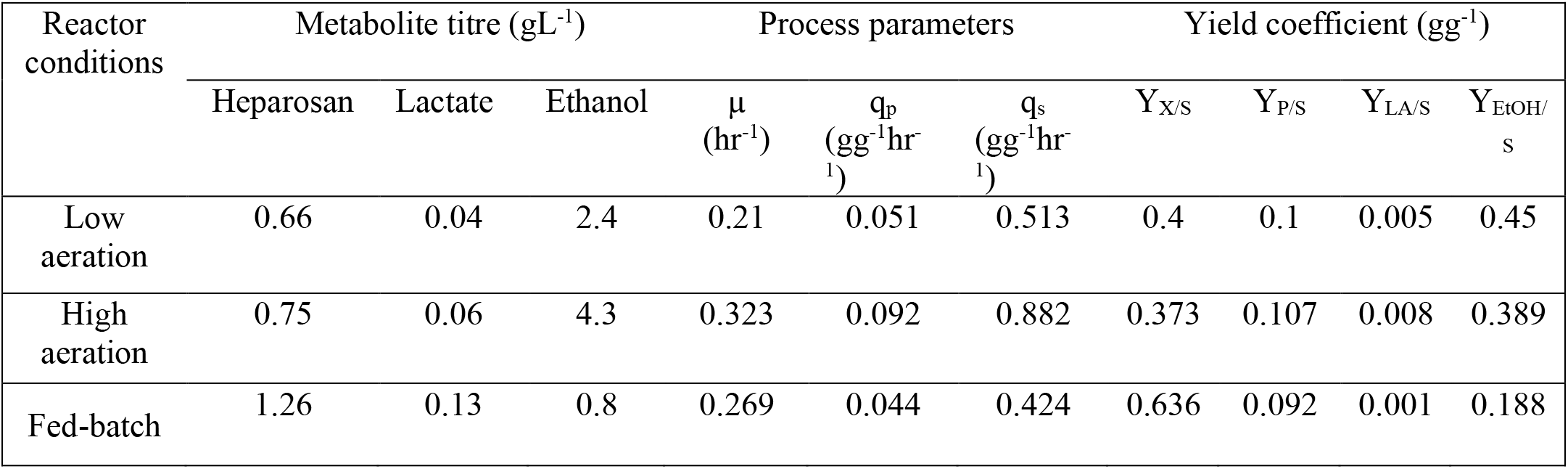
Comparison of metabolite titre, process parameters, and yield coefficients for SH6 clone, operated under different conditions in the fermenter.

This is because of the knockout of the gene Lactate dehydrogenase *ldh*, which is responsible for converting pyruvate to lactate. SH6 was found to have a product titre of 0.262 gL^-1^, the yield of product to the substrate Y_p/s_, and specific product formation rate q_p_ of 0.368 gg^-1^ and 0.059 gg^-1^hr^-1^. Lactate production was found to be low at 0.036 gL^-1^ only, which makes the downstream process much easier. The highest Y_p/s_ and q_p_ were observed in the SH6 clone, indicating that most of the carbon flux is passing through the heparosan-producing pathway. Owing to its low by-product formation, high Y_p/s_, q_p_ value, and good product titre, SH6 was chosen as the best and moved on to the reactor level.

### Microbial production of heparosan in fermenter

Batch fermentations were carried out using the clone SH6 at low aeration and high aeration with the initial glucose concentration of 30 gL^-1^. The kinetics of heparosan production in a batch fermenter under low aeration, high aeration, and fed-batch fermentations are depicted in Figure 2. It was observed that a batch reactor operating under high aeration conditions produces heparosan around 0.75 gL^-1^ and 0.66 gL^-1^ under low aeration conditions. Relatively higher product titre was observed under high aeration conditions than the low aeration conditions; this might be due to rerouting the carbon flux through NADH-independent pathways (Lopez de Felipe et al., 2006a). NAD+/NADH balance is required for maintaining the homeostasis of the cell, which could be achieved by the action of NADH oxidase under aerated conditions. Consequently, the carbon flux passes through the NADH-independent pathways resulting in higher heparosan production under high aeration conditions. A disaccharide unit (monomer) of Heparosan requires two glucose molecules, one acetyl coenzyme A, two UTP, and three ATP molecules, which is an energy-intensive process (Badle et al., 2014b). Under high aeration, the pyruvate produced via glycolysis might enter into the oxidative level phosphorylation, producing 32 net ATP molecules which might enhance the heparosan production. Hyaluronic acid, the heparosan analog, was likewise shown to have a similar effect, increasing product titre as the DO level increased (Jeeva et al., 2019). Because of the limiting substrate i.e. glucose exhaustion, the growth was hampered around the 20^th^ hour. Consequently, there was no heparosan production subsequently. To achieve high product titre, fed-batch fermentation was done by feeding the limiting substrate glucose into the fermenter once it was exhausted, which gave a higher heparosan of 1.26 gL^-1^. Despite being a growth-associated product, heparosan’s titre was found to drop after 30 hours as the biomass increased, as the same was also seen in (Zhang et al., 2012). This might be because of competition between heparosan synthesis and cell proliferation caused by glucose-1-P, a common precursor for both cell wall and heparosan synthesis (Zhang et al., 2012). The similar competition between cell growth and exopolysaccharide synthesis was also observed in hyaluronic acid production where glucose-1-phosphate, UDP-glucose, and UDP-N-acetylglucosamine act as a common precursor for both cell wall and hyaluronic acid synthesis (Yu & Stephanopoulos, 2008). This indicates a balance exists between cell growth and heparosan synthesis and that a limited amount of cell growth suppression may be advantageous for accumulating heparosan. It was also reported that in recombinant *E. coli* cultures, supplementation of fosfomycin (an inhibitor of cell wall synthesis) along with glucosamine reduced the growth and increased the hyaluronic acid (an analog of heparosan) synthesis (Mao et al., 2009).

**Figure 2:**
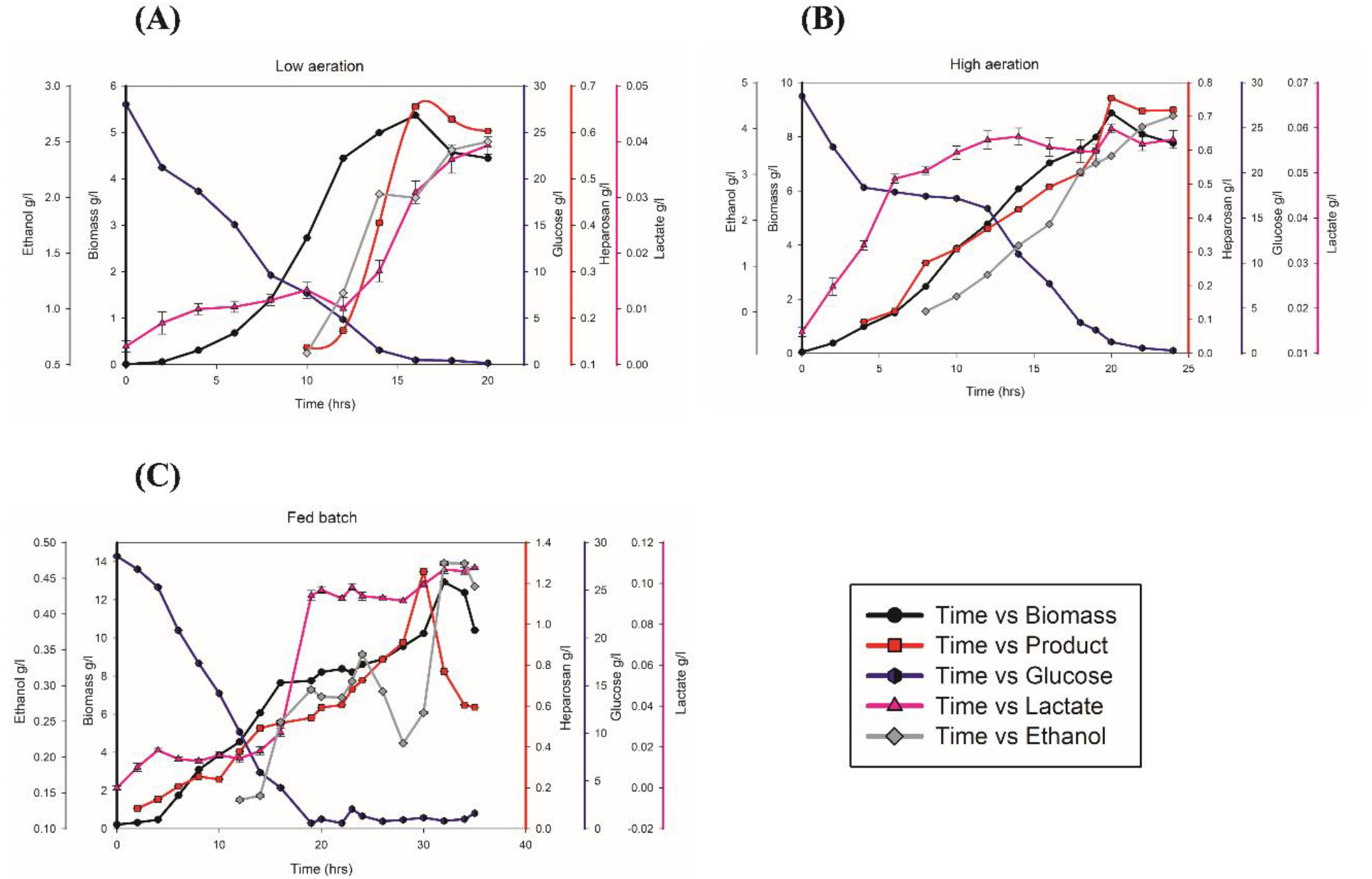
Kinetics of heparosan production in the bioreactor. Time profile of cell growth, heparosan concentration, Glucose consumption, lactate, and ethanol of SH6 in a bioreactor under (A) low aeration batch reactor, (B) high aeration batch reactor, (C) fed-batch reactor.

**Figure 3:**
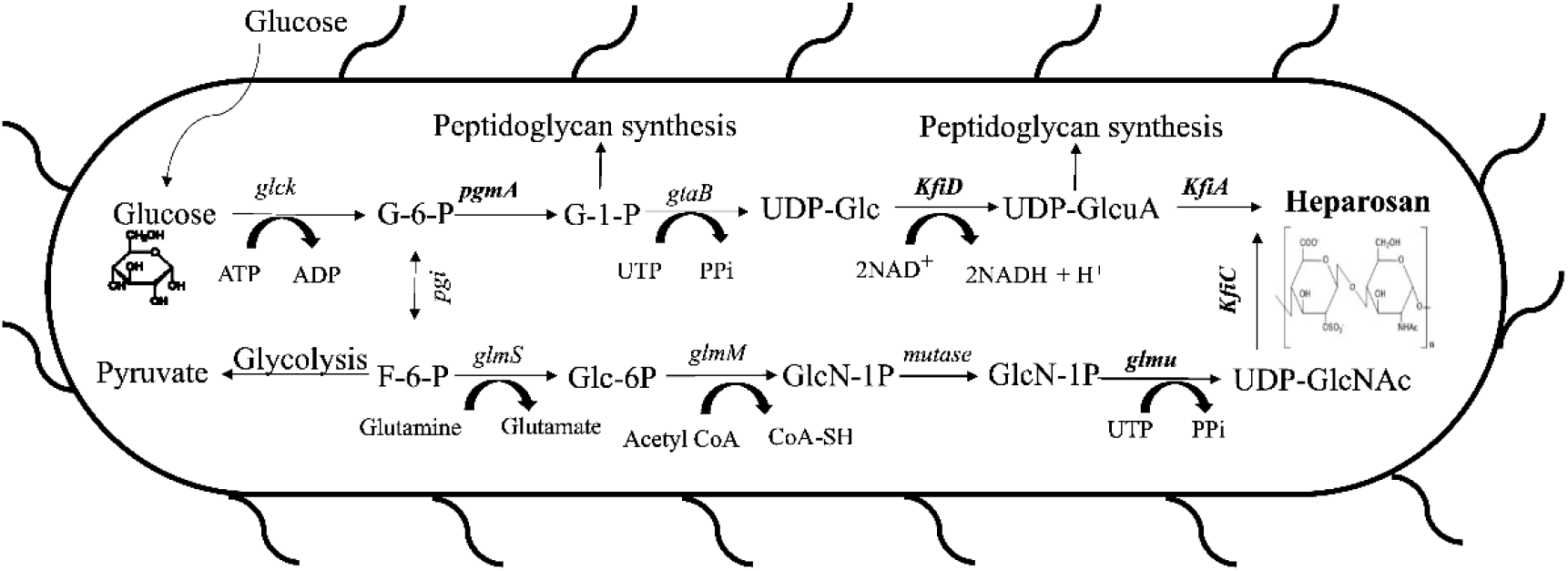
Heparosan biosynthesis pathway in recombinant *L. lactis*.

### Performance evaluation of SH6 clone under low aerated conditions

In the SH6 clone, lactate production was found to be very minimal, with 0.04 gL^-1^ having a yield coefficient Y_LA/S_ of 0.005 gg^-1^, which might be due to *ldh* gene knockout. *L. lactis* was generally found to be a strictly homofermentative LAB species, but it was observed that SH6 produces ethanol of 2.4 gL^-1^, in addition to heparosan. The yield of ethanol to the substrate Y_EtoH/S_ was found to be 0.45 gg^-1^. The balance of the NAD+/NADH ratio may be the primary cause of this. The NAD+/NADH ratio won’t be balanced by lactate production here for the SH6 clone because the *ldh* gene was knocked out. To compensate for this, pyruvate was transformed into acetate and, subsequently, ethanol by the action of alcohol dehydrogenase *aldh*, which produces NAD+, as seen in Figure 4. Hence the yield of ethanol with respect to the substrate was higher under low aeration conditions (Neves et al., 2005a). Also, it was observed that ethanol formation was found to be maximum when the glucose depletes below 5 gL^-1^. This indicates that *L. lactis* will produce ethanol under carbohydrate limitations (Pascal Hols et al., 1999). Additionally, it was stated that *L. lactis* would have increased glycolytic enzyme activity, producing an excess of pyruvate (Guillot et al., 2003a). To utilize the pyruvate under anaerobic conditions, ethanol formation might be happened, which was depicted in Figure 4.

**Figure 4:**
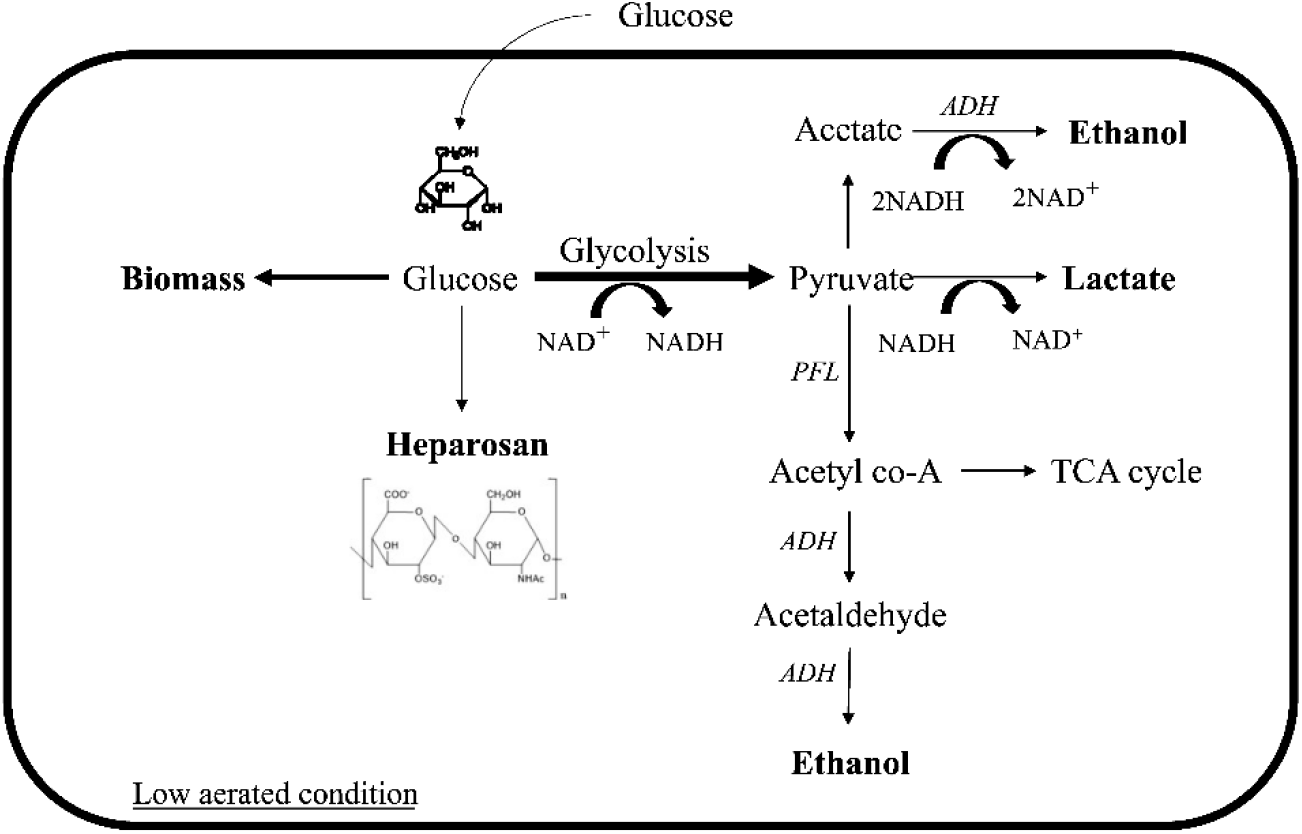
Carbon source utilization of recombinant *L. lactis* under low aerated batch conditions.

### Performance evaluation of SH6 clone under high aerated conditions

Under high aerated conditions, SH6 produces lactate and ethanol of 0.06 gL^-1^ and 4.3 gL^-1^ having yield coefficients Y_LA/S_, Y_EtoH/S_ of 0.008 gg^-1^, and 0.389 gg^-1^. Even though *L. lactis* is a homofermentative metabolism species that will convert sugars mainly to lactate; under certain conditions, such as carbohydrate limitation and aerobic fermentation, it will convert from homolactic fermentation to mixed-acid fermentation (Pascal Hols et al., 1999). *L. lactis* was reported to have higher glycolytic enzyme activity (Guillot et al., 2003b), resulting in the production of an enormous amount of pyruvate. H_2_O-forming NADH oxidase (NOX) got activated under aerobic conditions, which significantly reduced the amount of NADH available for *ldh;* as a result, pyruvate was redirected to the other aerobic catabolic pathways. Pyruvate will be converted to ethanol by the action of the pyruvate dehydrogenase complex (PDHC), which will be active under aerobic conditions (Lopez de Felipe et al., 2006b). Hence under aerobic conditions, the by-product ethanol formation was observed. Only at high quantities of NADH, the enzyme activity of PDHC will be inhibited. Since there won’t be much NADH due to NOX activity, there won’t be any inhibition of PDHC in aerobic conditions (Neves et al., 2005b). As a result, the high-aerated condition produces more ethanol than the low-aerated condition. Typically, *L. lactis* will produce lactate to keep the NAD+/NADH ratio balanced by converting NADH to NAD+. There is no need for lactate synthesis during aerobic conditions because NOX will convert NADH to NAD+. Also, due to the knockout of the gene *ldh*, which is primarily responsible for lactate generation, less lactate will be formed during the aerobic stages. Figure 5 depicts ethanol production and pyruvate metabolism under high aerated conditions.

**Figure 5:**
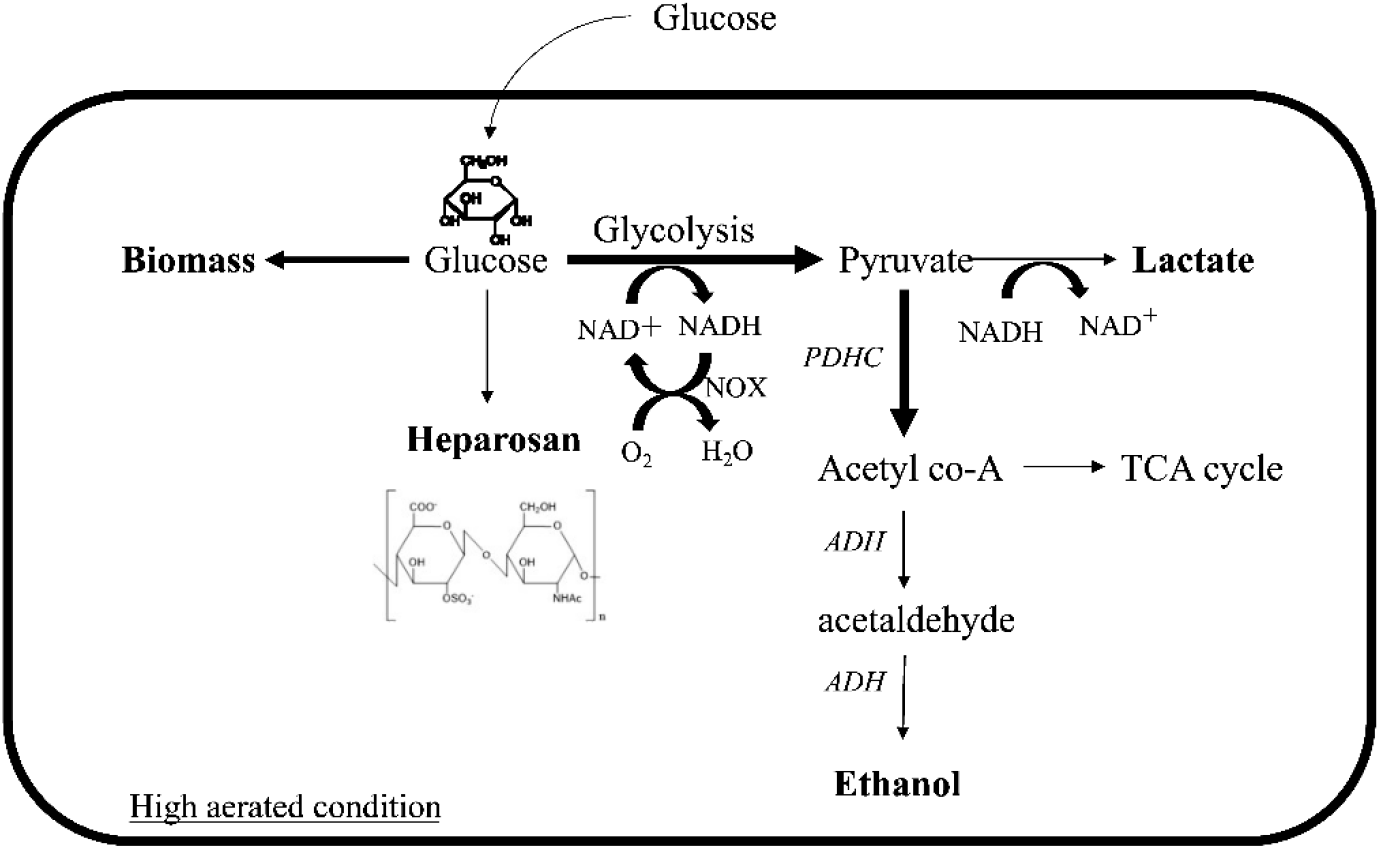
Carbon source utilization of recombinant *L. lactis* under high aerated batch conditions.

### Heparosan purification and molecular weight determination

The cell was lysed using lysis buffer and heat treatment, to release the intracellular heparosan. ATPS experiment revealed that almost all the heparosan was partitioned into the bottom phase, which was rich in salt. Heparosan was concentrated via aqueous two-phase extraction at the salt-rich bottom phase, with a partition coefficient of 0.22 and a 78% recovery rate. Heparosan settles to the bottom phase as a result of the steric exclusion of polymer PEG 6000 in the top phase. To get rid of the potassium phosphate salt, dialysis was done.

Molecular weight is one of the critical parameters that need to be considered for heparin production. The heparosan molecule purified yielded a retention time of 19.2 min on the GPC column, giving a molecular weight of around 10-20 kDa. The molecular weight range of 10 kDa to 20 kDa produced by *L. lactis* engineered with *E. coli K5* glycosyltransferases highlights its potential as a precursor for producing heparin. Low molecular heparin of around 12 kDa is mainly used in the pharmaceutical industry. Because of their higher subcutaneous bioavailability and superior pharmacokinetic/pharmacodynamic characteristics, Low Molecular Weight Heparosan (LMWH) has a significant advantage in biological half-lives (Linhardt, 2003). Cellular doubling time and specific growth rate are inversely correlated. Higher doubling times allow the cell more time to elongate the heparosan chain before dividing. It was reported that the glycosyl transferase polymerization rate for increasing the chain length is around 20 to 10,000 monomers per hour (Deangelis, 2002). Since the doubling time of *L. lactis* is comparable to the rate of polymerization of a glycosyl transferase, the specific growth rate may play an essential role in the heparosan chain termination and lead to the production of low molecular weight heparosan. The competition between precursors may impact the polymerization activity of glycosyl transferase due to the affinity of UDP-GlcUA to the UDP-GlcNAc binding site or vice versa. When one of the sugar nucleotides binds to both sites simultaneously, chain termination may result. This may occur if one of the precursors has a significantly higher concentration than the other. Because *ugd* and *pgma*, the genes responsible for producing UDP-GlcUA, were overexpressed in the SH6 clone. Chain termination and the generation of low molecular weight heparosan may arise from this. Hence it was observed that the product titre and the molecular weight were inversely correlated.

### Chemical structure characterization of Heparosan from *L. lactis* using FTIR and NMR

To determine the structure of heparosan obtained from *L. lactis*, FTIR and one-dimensional ^1^H NMR and ^13^C NMR investigations were carried out. FTIR spectra obtained for the clone SH6 are shown in Figure 6A, which correlates well with the articles that have been previously published (Lokwani et al., 2015; Qiao et al., 2009), and their corresponding functional group peak ranges are shown in Table 6. ^1^H NMR and ^13^C NMR were used to describe the heparosan that was produced from the clone SH6. The chemical shifts of the proton and carbon are shown in Table 7, and they are strongly supported by the heparosan spectra that have already been published (Datta et al., n.d.; Nehru et al., 2020; Zhang et al., 2012). The peaks in Figure 6B with asterisks are the molecular characteristics and are connected to the C-3, C-4, and C-5 of GlcUA and GlcNAc. These findings suggest that the structure of heparosan produced from recombinant *L. lactis* has identical disaccharide repeating units.

**Table 6:** Comparison of FTIR frequencies of *L. lactis*-derived heparosan with previous studies

| Functional groups | Wavenumber (cm<sup>-1</sup>) | Reported range |
| --- | --- | --- |
| O-H groups stretch | 3412 | 3100-3700 (Qiao et al., 2009) |
| N-H bond stretch | 1624 | 1600-1700 (Qiao et al., 2009) |
| H-C-H bond stretch | 2434 | 2400-2600 (Lokwani et al., 2015) |

**Table 7:**
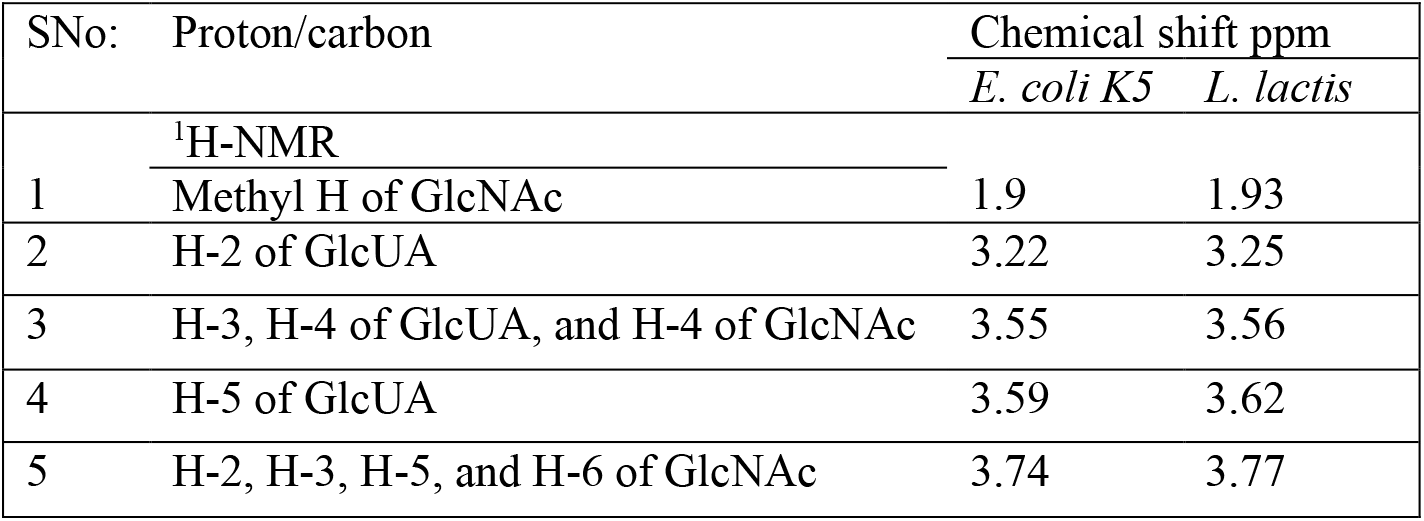
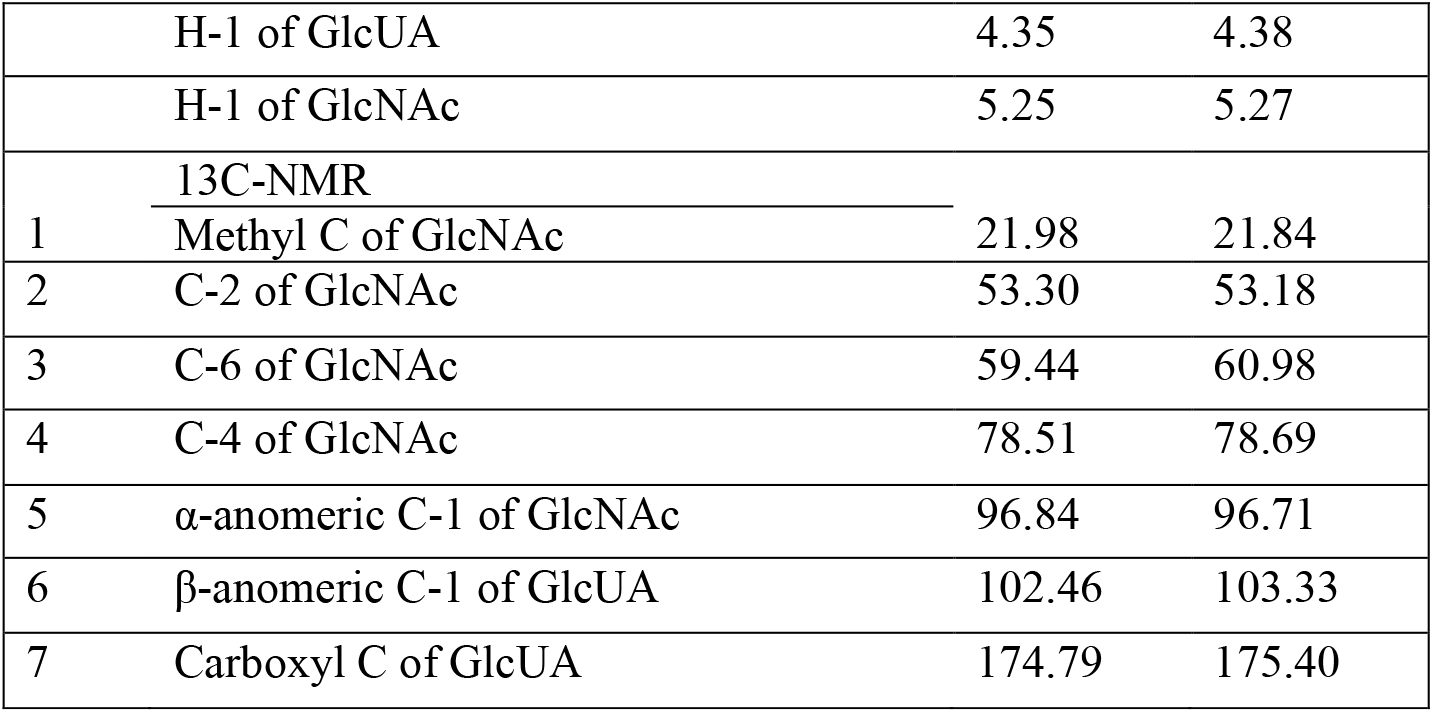
Chemical shift assignments of *L. lactis-derived* heparosan. Chemical shifts are represented in ppm.

| SNo: | Proton/carbon | Chemical shift ppm |  |
| --- | --- | --- | --- |
|  |  | *E. coli* K5 | *L. lactis* |
|  | <sup>1</sup>H-NMR |  |  |
| 1 | Methyl H of GlcNAc | 1.9 | 1.93 |
| 2 | H-2 of GlcUA | 3.22 | 3.25 |
| 3 | H-3, H-4 of GlcUA, and H-4 of GlcNAc | 3.55 | 3.56 |
| 4 | H-5 of GlcUA | 3.59 | 3.62 |
| 5 | H-2, H-3, H-5, and H-6 of GlcNAc | 3.74 | 3.77 |
|  | H-1 of GlcUA | 4.35 | 4.38 |
|  | H-1 of GlcNAc | 5.25 | 5.27 |
|  | <u><sup>13</sup>C-NMR</u> |  |  |
| 1 | Methyl C of GlcNAc | 21.98 | 21.84 |
| 2 | C-2 of GlcNAc | 53.30 | 53.18 |
| 3 | C-6 of GlcNAc | 59.44 | 60.98 |
| 4 | C-4 of GlcNAc | 78.51 | 78.69 |
| 5 | α-anomeric C-1 of GlcNAc | 96.84 | 96.71 |
| 6 | β-anomeric C-1 of GlcUA | 102.46 | 103.33 |
| 7 | Carboxyl C of GlcUA | 174.79 | 175.40 |

**Figure 6:**
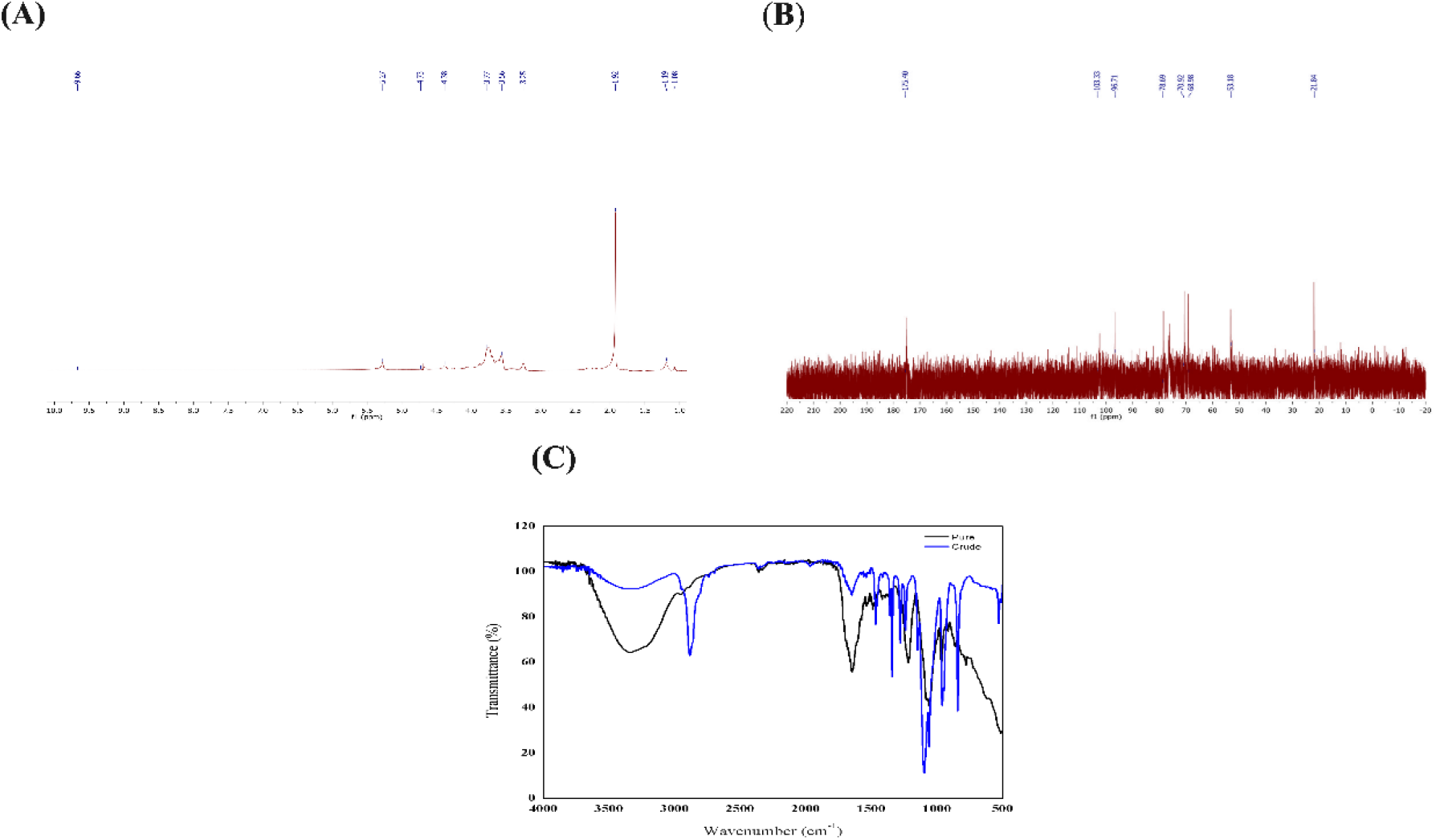
(A) 1H-NMR and (B) 13C-NMR spectra of heparosan from recombinant *L. lactis*. (C) FTIR spectra of heparosan from recombinant L. lactis and their corresponding wavenumbers.

## Conclusion

This study successfully addressed a safe and alternative method to generate heparosan in the GRAS micro-organism *L. lactis* using *E. coli K5* glycosyltransferase genes. The yield of six distinct clones with different gene combinations for the production of the intracellular precursor for heparosan was analyzed. Heparosan concentrations reached a maximum of 262 mgL^-1^ during batch fermentation and 1.26 gL^-1^ during fed-batch fermentation. Heparosan that was produced in the molecular weight range (10–20 kDa) could serve as a precursor to chemoenzymatic heparin production. Aeration rate, agitation speed, and dissolved oxygen content, in addition to the media, may also significantly impact the production of heparosan by *L. lactis*. These process optimizations can be carried out to further improve the titre of heparosan in the future.

## Notes

### Competing Interest Statement

The authors have declared no competing interest.

